# Non-viral induction of transient cell reprogramming in skeletal muscle to enhance tissue regeneration

**DOI:** 10.1101/101188

**Authors:** Irene de Lázaro, Acelya Yilmazer, Yein Nam, Sarah Qubisi, Fazilah Maizatul Abdul Razak, Giulio Cossu, Kostas Kostarelos

## Abstract

Somatic cells can be reprogrammed to pluripotency *in vivo* by overexpression of defined transcription factors. While their sustained expression triggers tumorigenesis, transient reprogramming induces pluripotency-like features and proliferation only temporarily, without teratoma formation. We sought to achieve transient reprogramming within mouse skeletal muscle with a localized injection of plasmid DNA (pDNA) and hypothesized that this would enhance regeneration after severe injury. Intramuscular administration of reprogramming pDNA rapidly upregulated pluripotency (*Nanog*, *Ecat1*, *Rex1*) and early myogenesis genes (*Pax3*) in the healthy gastrocnemius of various mouse strains. Mononucleated cells expressing such markers appeared promptly in clusters among myofibers, but proliferated only transiently and did not lead to the generation of teratomas. *Nanog* was also upregulated in the gastrocnemius when reprogramming factors were administered 7 days after laceration of its medial head. Enhanced tissue regeneration after reprogramming was manifested by the accelerated appearance of centro-nucleated myofibers and reduced fibrosis. These results suggest that *in vivo* transient reprogramming may constitute a novel strategy towards the acceleration of regeneration following muscle injury, based on the induction of transiently-proliferative, pluripotent-like cells *in situ*. Further research to achieve clinically meaningful functional regeneration is warranted.

## Introduction

Forced expression of reprogramming transcription factors, among them the combination known as ‘Yamanaka’ or ‘OKSM’ factors (Oct3/4, Klf4, Sox2 and cMyc), results in the reversion of differentiated cells to a pluripotent state, not only in the culture dish [1] but also within adult tissues *in vivo* [2-7]. Such *in vivo* conversion to pluripotency has now been described in a variety of transgenic and wild-type (WT) animal models in spite of the pro-differentiation signals present in the tissue microenvironment. Ubiquitous and/or sustained expression of reprogramming factors *in vivo* leads to uncontrolled proliferation and widespread tumorigenesis [4-7]. Conversely, studies have shown that transient expression of such genes results in the generation of pluripotent intermediates that proliferate only temporarily and do not produce dysplastic lesions or teratomas in the tissue [2, 3, 5]. We have previously shown this effect in WT mouse liver by using a nonviral approach based on hydrodynamic tail vein (HTV) injection of naked plasmid DNA (pDNA) encoding the Yamanaka factors[3, 8]. When isolated from the liver, *in vivo* reprogrammed cells generated cells and tissues from all three germ layers in a variety of *in vitro* and *in vivo* assays, which confirmed their functional pluripotency [9].

In the adult mammalian organism, muscle repair relies mainly on the self-renewal and myogenic potential of resident stem cells, primarily satellite cells [10]. The number and regenerative capacity of such cells varies across species and dramatically decreases with age [11, 12], hence it can be easily exhausted if the injury is severe or repeated. In this scenario, various resident cell types differentiate into myofibroblasts that generate a collagen-based fibrotic scar unable to meet the contractile requirements of the tissue, therefore preventing the complete functional rehabilitation of the injured muscle [13, 14].

Different strategies are currently being explored to enhance muscle regeneration after severe injuries, including surgical suturing [15] or administration of anti-fibrotic drugs [13, 14, 16-18], growth factors [19], replacement cells [20-22], combinations of these [23-25] and miRNAs involved in muscle development [26]. However, none of these approaches has yet reached routine clinical practice and the treatment and management of major muscle injuries continues to be primarily conservative [27, 28]. At the same time, the skeletal muscle tissue offers an excellent platform for the expression of foreign genes *in vivo*, given the capacity of the myofibers to uptake naked pDNA after a simple intramuscular (i.m.) administration[29]. Although the exact mechanism by which myofibers incorporate the naked nucleic acid has not yet been fully elucidated, working hypotheses include an active uptake mechanism [30] and the use of the T tubule system [31]. In addition, pDNA is also taken up by interstitial mononucleated cells located between myofibers upon intramuscular administration [32].

We have previously hypothesized that the generation of pluripotent or pluripotent-like cells *in vivo* able to transiently proliferate and eventually re-differentiate to the appropriate phenotype within a damaged tissue may assist its cellular repopulation and therefore boost its capacity to regenerate after a traumatic insult or injury[33]. In the present work, we aimed first to explore whether transient, teratoma-free pluripotent or pluripotent-like conversion could be achieved in mouse skeletal muscle via i.m. administration of naked reprogramming pDNA, to then interrogate whether such strategy would enhance the regenerative capacity of the muscle after a surgically-induced injury.

## Results

### Gene and protein expression in mouse skeletal muscle after i.m. administration of reprogramming factors

To test whether pluripotency could be induced in mouse skeletal muscle, we first i.m. administered BALB/c mice with 50 μg of a single pDNA cassette encoding OKSM factors and an eGFP reporter (pLenti-lll-EF1a-mYamanaka) in the gastrocnemius (GA) muscle. The contralateral hind limb was injected with the same volume of saline solution and used as control. Changes in gene and protein expression in the GA muscle were investigated at different time points after injection by real-time RT-qPCR and immunohistochemistry (IHC), respectively (Figure 1). 2 days after injection, the reprogramming factors (*Oct3/4*, *Sox2*, *c-Myc*) and *eGFP* reporter were highly expressed (mRNA level) in the muscles administered with reprogramming pDNA to then decline and, in the case of *c-Myc* and *eGFP*, reach baseline levels as early as 4 days after *in vivo* transfection (Figure 1b). This rapid decrease in the expression of the transgenes contrasted with the stable and long-term foreign gene expression that is normally observed after the uptake of pDNA in post-mitotic myofibers [34]. In agreement with this, eGFP localized mainly in interstitial mononucleated cells (Figure 1f).

**Figure 1.**
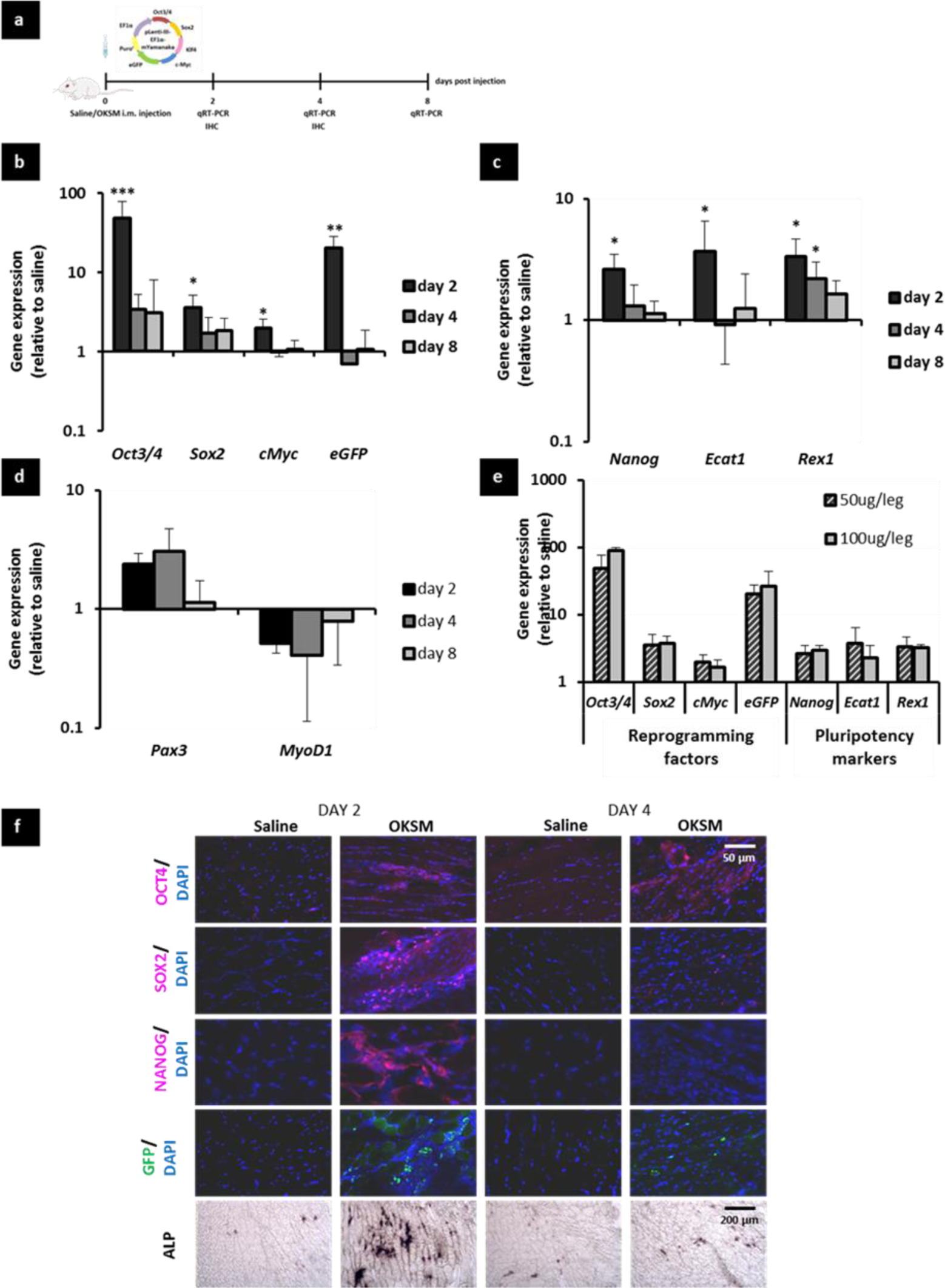
Gene and protein expression in the mouse skeletal muscle after i.m. administration with reprogramming pDNA. **(a)**BALB/c mice were i.m. injected in the GA muscle with 50 μg pLenti-lll-EF1a-mYamanaka in 50 μl 0.9% saline or 50 μl saline alone. Ga muscles were dissected 2, 4 and 8 days after the injection and real-time RT-qPCR was performed to determine the relative gene expression of **(b)** the transfected reprogramming factors, **(c)** endogenous pluripotency markers and **(d)** genes involved in myogenesis.**(e)** Dose response was obtained with 50 and 100μg pLenti-lll-EF1a-mYamanaka per GA muscle. Gene expression levels were normalised to the saline-injected group. ***p<0.001. **p<0.01 and *p<0.05 indicate statistically significant differences in gene expression between pDNA and saline-injected groups, assessed by one-way ANOVA or Welch ANOVA. Data are presented as mean ± SD, n=3. **(f)** Frozen tissue section from Ga muscles dissected on days 2 and 4 were stained with anti-OCT4, anti-SOX2 or anti-NANOG antibodies to assess immunoreactivity, or BCIP/NBT to determine AP activity, or directly observed under an epifluorescence microscope to detect the expression of the eGFP reporter in the tissue. Fluorescence images were taken at 40X magnification, scale bars represent 50μm. Brightfield images were taken at 10X magnification, scale bars represent 200μm

Five-fold upregulation of genes endogenously expressed in the pluripotent state but typically repressed in adult tissues (*Nanog*, *Ecat1*, *Rex1*) was also observed as early as day 2 and decreased afterwards (Figure 1c). The investigation of genes characteristically expressed at different stages of myogenesis indicated the upregulation of *Pax3,* that is distinctly expressed in myogenic progenitors, but not in differentiated myofibers [35], only for the OKSM injected group. On the contrary, *MyoD1*, that is only expressed in more committed stages during skeletal muscle development [36] and is a direct target of *Oct4* during myoblast-to-iPS cell reprogramming – hence should be downregulated during such process [37] – was indeed found to be expressed at lower levels compared to saline-injected controls (Figure 1d). These results suggested the activation of an embryonic-like gene expression program in a subset of cells within the skeletal muscle upon forced expression of reprogramming factors *in vivo*. The administration of a higher dose (100 μg) of pLenti-lll-EF1a-mYamanaka did not result in significant changes in the expression of reprogramming or pluripotency markers (Figure 1e) and hence the lowest dose of reprogramming pDNA (50 μg) was selected for further studies.

The induction of pluripotent-like features in the tissue was further confirmed by IHC (Figure 1f). Clusters of mononucleated cells expressing pluripotency markers (OCT4, SOX2 and NANOG) and exhibiting high alkaline phosphatase (AP) activity – a recognized feature of pluripotent cells [38] – were observed among the myofibers 2 days after injection. The expression of NANOG was cytoplasmic, instead of its most common localization in the cell nucleus. However, such event has been previously described by others [39, 40]. Green fluorescence resulting from the eGFP reporter in pLenti-lll-EF1a-mYamanaka was bright in the clusters, confirming that such cells had been transfected with reprogramming pDNA. Immunostaining for the above markers was not detected from day 4 after i.m. injection, which supported our observations at the mRNA level and confirmed that the conversion to a pluripotent-like state occurred very rapidly, yet transiently.

### *In vivo* reprogramming towards pluripotency in Nanog-GFP and Pax3-GFP transgenic mice skeletal muscle

Sv129-Tg(Nanog-GFP) transgenic mice, referred to as Nanog-GFP for simplicity here, have the reporter GFP sequence inserted in the *Nanog* locus [41] and therefore were used to identify cells exhibiting pluripotency features in the tissue thanks to the emission of green fluorescence. pCX-OKS-2A and pCX-cMyc cassettes were used instead of pLenti-lll-EF1a-mYamanaka to avoid overlap with the eGFP reporter encoded in the latter (Figure 2a). Prior to any histological analysis, we confirmed that the expression of reprogramming factors, endogenous pluripotency genes and myogenesis-related markers on days 2, 4 and 8 after i.m. injection was comparable to that observed in WT mice (Figure 2b-d).

**Figure 2.**
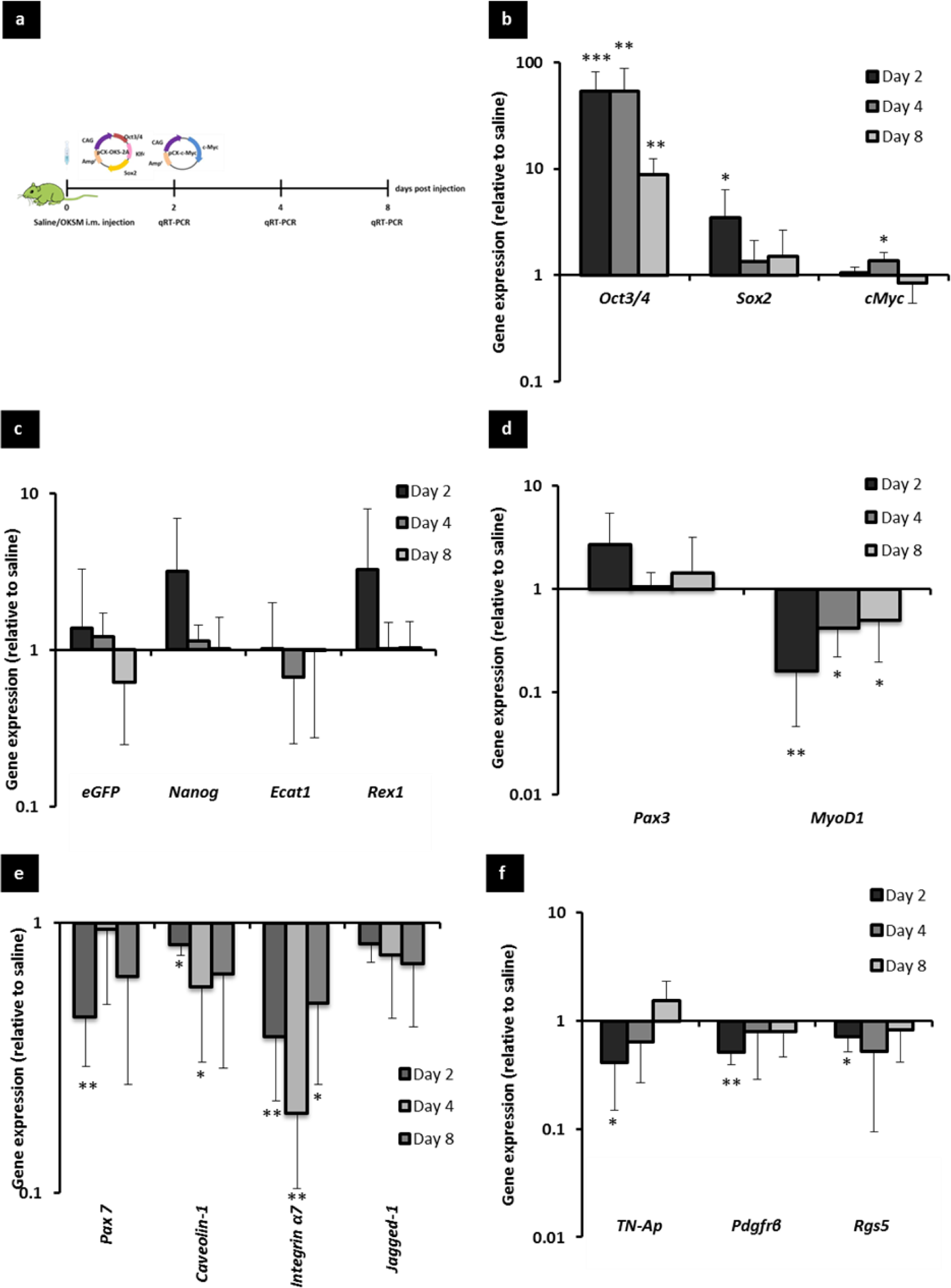
Gene expression in the Nanog-GFP transgenic mouse skeletal muscle after i.m. administration with reprogramming pDNA. **(a)** Nanog-GFP mice were i.m. injected in the GA muscle with 50 μg pCX-OKS-2A and 50 μg pCX-cMyc in 50 μl 0.9% saline or 50 μl saline alone. GA muscles were dissected 2, 4 and 8 days after the injection and real-time RT-qPCR was performed to determine the relative gene expression of **(b)** the transfected reprogramming factors, **(c)** endogenous pluripotency markers, **(d)** genes involved in myogenesis.**(e)** satellite cell markers and **(f)** pericyte markers. Gene expression levels were normalised to the saline-injected group. *p<0.05, **p<0.01 and ***p<0.001 indicate statistically significant differences in gene expression between pDNA and saline-injected groups, assessed by one-way ANOVA or Welch ANOVA. Data are presented as mean ± SD, n=4.

While the observations in this mouse strain also suggested the reprogramming of a subset of cells towards an embryonic-like state, the input of pericytes and satellite cells that are present in the skeletal muscle and have myogenic potential needed to be clarified [10, 42]. Satellite cells express *Pax3* in certain muscles[35]. Therefore, it was of particular importance to rule out the contribution of a hypothetical satellite cell proliferation derived from the injection trauma towards the elevated mRNA levels of such transcription factor. The expression of satellite cell-specific markers (*Pax7*, *Caveolin1*, *Integrin-α7*, *Jaggedl*) and pericyte markers (*TN-AP*, *Pdgfrβ, Rgs5*) was hence investigated and found to be downregulated compared to controls (Figure 2 e-f).

At the histological level (Figure 3), H&E staining revealed the presence of dense clusters of mononucleated cells among the myofibers in Nanog-GFP mice administered with OKSM. The bright green fluorescence signal observed 2 days after injection (Figure 3b) that confirmed the expression of *Nanog* and hence the pluripotent-like identity of such clusters, was not detected at later time points (data not shown). This finding further suggested the transiency of the conversion to such pluripotent-like state. The GFP signal co-localised with the expression of several other pluripotency and embryonic stem cell specific markers identified by IHC (NANOG, OCT4, AP and SSEA1) (Figure 3c). No GFP signal or immunoreactivity for any of the pluripotency markers tested was found in saline-injected controls, excluding AP, which is also expressed by pericytes (**Figure S1b and d**).

**Figure 3.**
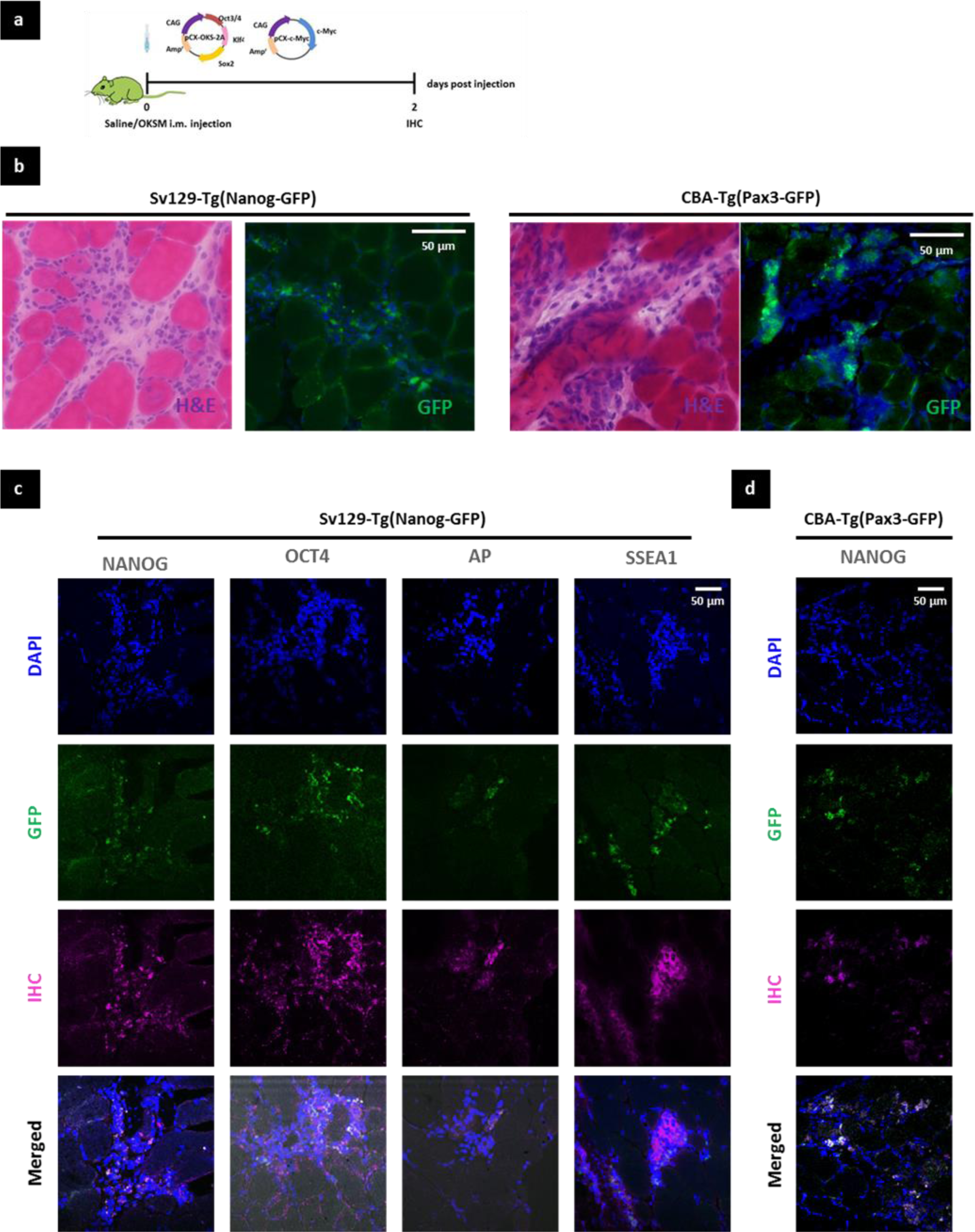
Characterisation of *in vivo* reprogrammed cell clusters in the GA muscle of Nanog-GFP and Pax3-GFP transgenic mice. **(a)** Nanog-GFP and Pax3-GFP mice were i.m. administered with 50 μg pCX-OKS-2A and 50 μg pCX-cMyc in 50 μl 0.9% saline or 50 μl saline alone. GA muscles were dissected 2 days after injection and 10 μm-thick tissue sections were obtained by cryotomy. (**b**) Clusters of reprogrammed cells in the GA muscle were identified by H&E and the GFP signal resulting from either *Nanog* or *Pax3* upregulation (100X, scale bars represent 50 μm). Brightfield and fluorescence images show the same region within the tissue. (**c**) IHC for the expression of pluripotency markers in Nanog-GFP mouse muscle tissue administered with reprogramming pDNA, (100X, scale bars represent 50 μm). (**d**) IHC for the expression of the pluripotency marker NANOG in Pax3-GFP mouse muscle tissue administered with reprogramming pDNA, (100X, scale bars represent 50 μm).

We also found cells staining positively for satellite cell and pericyte markers (PAX7 and PDGFrβ, respectively) in the tissues from pDNA-injected mice (**Figure S1c**). However, such cells were found in similar numbers as in saline-injected controls (**Figure S1d**) and were located in the vicinity of the clusters, but not co-localizing with the GFP^+^ cells within them. This finding reaffirmed our observations at the mRNA level (Figure 2e-f) and indicated that the events triggered in the skeletal muscle upon *in vivo* reprogramming towards pluripotency were different from a proliferative response of resident stem/progenitor cells.

The use of a CBA-Tg(Pax3-GFP) transgenic mouse strain, referred to as Pax3-GFP, also allowed the identification of bright GFP^+^ cell clusters 2 days after pDNA injection, which were very similar to those observed in Nanog-GFP mouse tissues and did not appear in the saline-injected controls (Figures 3b **and S1b**). This observation agreed with our gene expression data that indicated the upregulation of *Nanog* and *Pax3* on day 2 after injection (Figure 2c-d), but did not confirm whether both markers were co-expressed by the same cells. To answer this, we performed anti-NANOG IHC on Pax3-GFP mouse skeletal muscle tissue sections and found precise co-localisation of both signals in cells within the clusters (Figure 3d). This finding confirmed that reprogrammed cells expressing NANOG also expressed PAX3.

Overall, these data suggested that the cells reprogrammed to a pluripotent-like state within the skeletal muscle tissue grew in clusters of mononucleated cells and expressed pluripotency and early myogenic progenitor markers, but not those characteristic to satellite cells or pericytes. While not all the cells within the clusters in Nanog-GFP and Pax3-GFP specimens expressed the green reporter, this heterogeneity might also be explained by the transiency of the reprogramming event, with some cells probably re-differentiating already at the time point investigated (day 2). The occurrence of partial reprogramming events cannot also be ruled out.

### Short and long-term effects of *in vivo* reprogramming towards pluripotency in healthy skeletal muscle

We next focused on the outcome of the forced expression of reprogramming factors (pLenti-lll-EF1a-mYamanaka) in healthy BALB/c GA with an emphasis on signs of regeneration, cell proliferation and potential appearance of dysplastic lesions or teratomas. Both short and long time points after the induction of pluripotency - from day 2 to 120 after injection - were studied (Figure 4a). In agreement with the observations in Nanog-GFP and Pax3-GFP mice, H&E and desmin/laminin stainings evidenced the appearance of distinct clusters of mononucleated cells in the OKSM-administered group, which were very prominent and densely packed 2 days after injection (Figure 4b-c). From day 4, small calibre, desminpositive, centronucleated myofibers (characteristic features of regenerating, immature myofibers[43, 44]) appeared within such clusters. Only few desmin-positive, centronucleated myofibers were noted on day 8 after injection and none at later time points. Lower magnification images of these observations are shown in **Figure S2b**.

**Figure 4.**
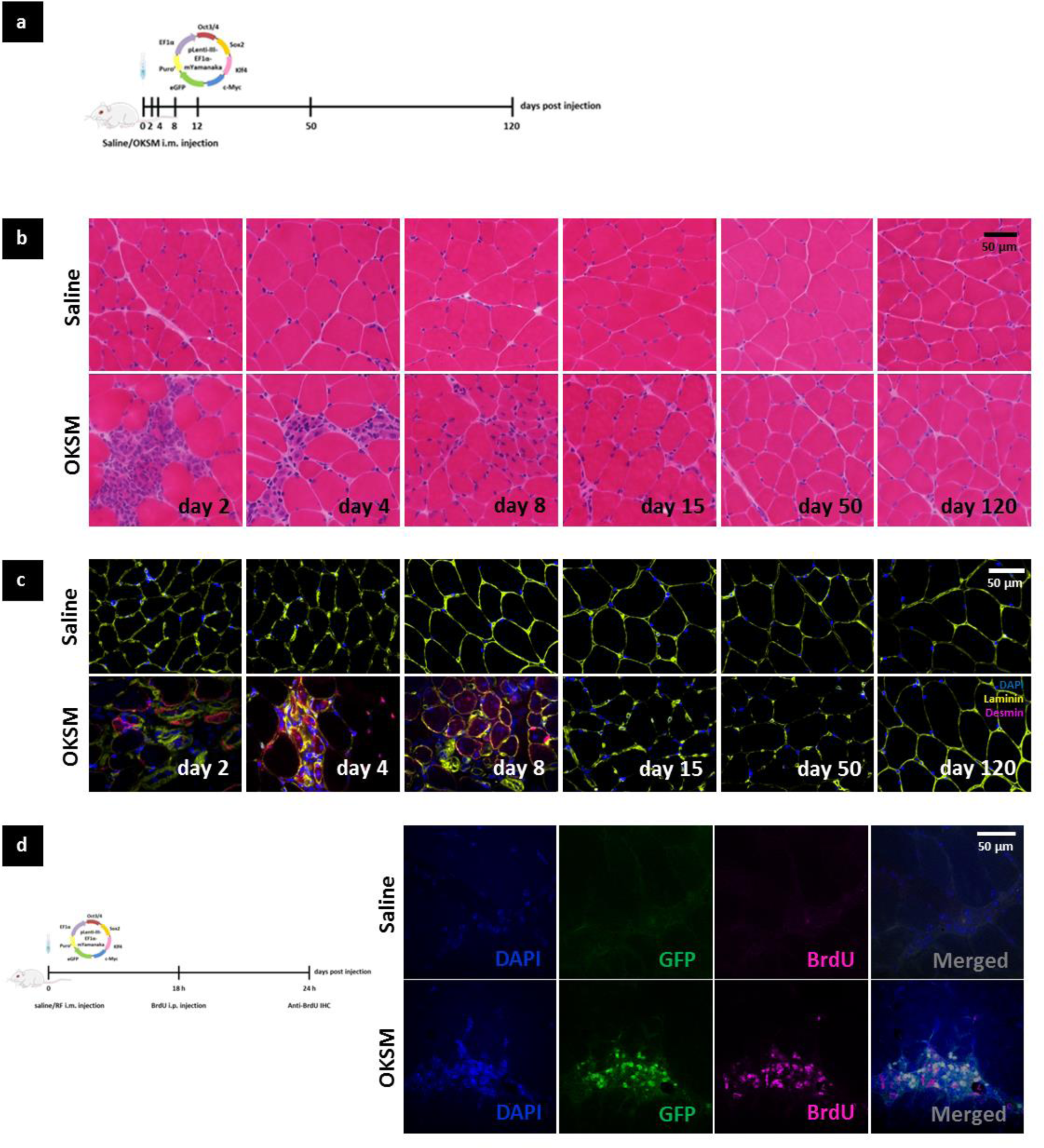
The effect of *in vivo* reprogramming towards pluripotency on healthy skeletal muscle tissue. **(a)** BALB/c mice were i.m. injected in the GA muscle with 50μg pLenti-lll-EF1a-mYamanaka in 50 μl 0.9% saline or 50 μl saline alone. GA muscles were dissected 2, 4, 8, 12, 50 and 120 days after injection.10 μm-thick tissue sections obtained by cryotomy were stained with (**b**) H&E (100X, scale bar represents 50 μm) and (**c**) anti-desmin and anti-laminin antibodies (40X, scale bar represents 50 μm). (**d**) BALB/c mice were i.p. injected with 500mg/kg BrdU 18 h after the i.m. administration of 50μg pLenti-lll-EF1a-mYamanaka in 50 μl 0.9% saline or 50 μl saline alone in the GA muscle. GA muscles were dissected 24 h after pDNA injection. 10 μm-thick tissue sections obtained by cryotomy were stained with anti-BrdU antibody. GFP signal corresponded to the reporter encoded in the pDNA. Images were captured with a confocal microscope (100X). Scale bar represents 50 μm.

We then sought to obtain more stringent evidence of cell proliferation that would reaffirm our hypothesis. Cell division has indeed been identified as a distinct and indispensable step in the somatic-to-pluripotent conversion[45, 46]. To address this, 18 h after i.m. administration with 50 μl of pLenti-lll-EF1a-mYamanaka or saline control, BALB/c mice were i.p. injected with 5-Bromo-2’deoxyuridine (BrdU), which can be incorporated in the DNA of proliferating cells[47]. Tissues were collected 6 h later to undertake histological examination (Figure 4d). GA muscle tissue sections were IHC stained with an anti-BrdU antibody and the eGFP reporter in pLenti-lll-EF1a-mYamanaka was used to identify the cells transfected with reprogramming factors. We found co-localisation of the two signals in cell clusters within the GA tissue in OKSM-injected animals, which were very similar to those observed in our previous studies. This finding confirmed that *in vivo* reprogrammed cells not only acquired a pluripotent-like gene expression profile, but also proliferated actively. However, the progressive disappearance of cell clusters together with the absolute lack of teratoma formation and/or dysplastic abnormalities observed for the duration of the study suggested that such proliferative state was only transient (Figure 4b).

In addition, no differences were observed in the abundance of apoptotic nuclei between the OKSM and saline-injected groups. TUNEL staining indicated the presence of limited numbers of apoptotic nuclei only at the earliest time points after injection and in both conditions; hence their occurrence was attributed to mild tissue damage caused by the injection along the needle track (**Figure S2c**). This suggested that the *in vivo* reprogrammed cells successfully re-integrated into the muscle tissue, although appropriate lineage tracing models will be necessary to ultimately clarify their fate.

### Enhancement of regeneration in a surgically-induced model of skeletal muscle injury

The capacity of *in vivo* reprogramming towards pluripotency to enhance regeneration was interrogated in a clinically relevant model of skeletal muscle injury. The left medial GA muscle of BALB/c mice was surgically lacerated in the transverse plane as represented in **Figure S3a** and pLenti-lll-EF1a-mYamanaka was i.m. administered in the injured hind limb at the time of injury, 5 or 7 days later. Control animals bearing the same injury were injected with 0.9% saline solution alone and the contralateral (right) leg of each mouse was left intact (uninjured and uninjected) as internal control. Analysis of *Nanog* mRNA levels in the tissue 2 days after the administration of reprogramming pDNA indicated that maximum upregulation of the pluripotency marker was achieved when the Yamanaka factors were administered 7 days after injury (**Figure S3b**). Muscle regeneration was then studied 9 and 14 days after injury (2 and 7 days after administration of reprogramming factors or saline control) (Figure 5a). The uptake and expression of the reprogramming pDNA, evidenced by *Oct3/4* expression, was confirmed on day 9. mRNA levels of the transgene were significantly higher in the lateral GA compared to the directly injured medial GA, possibly due to the abundance of necrotic myofibers at the injury site. Likewise, the upregulation of the pluripotency marker *Nanog* was also more pronounced in the lateral head of the GA muscle (Figure 5b).

**Figure 5.**
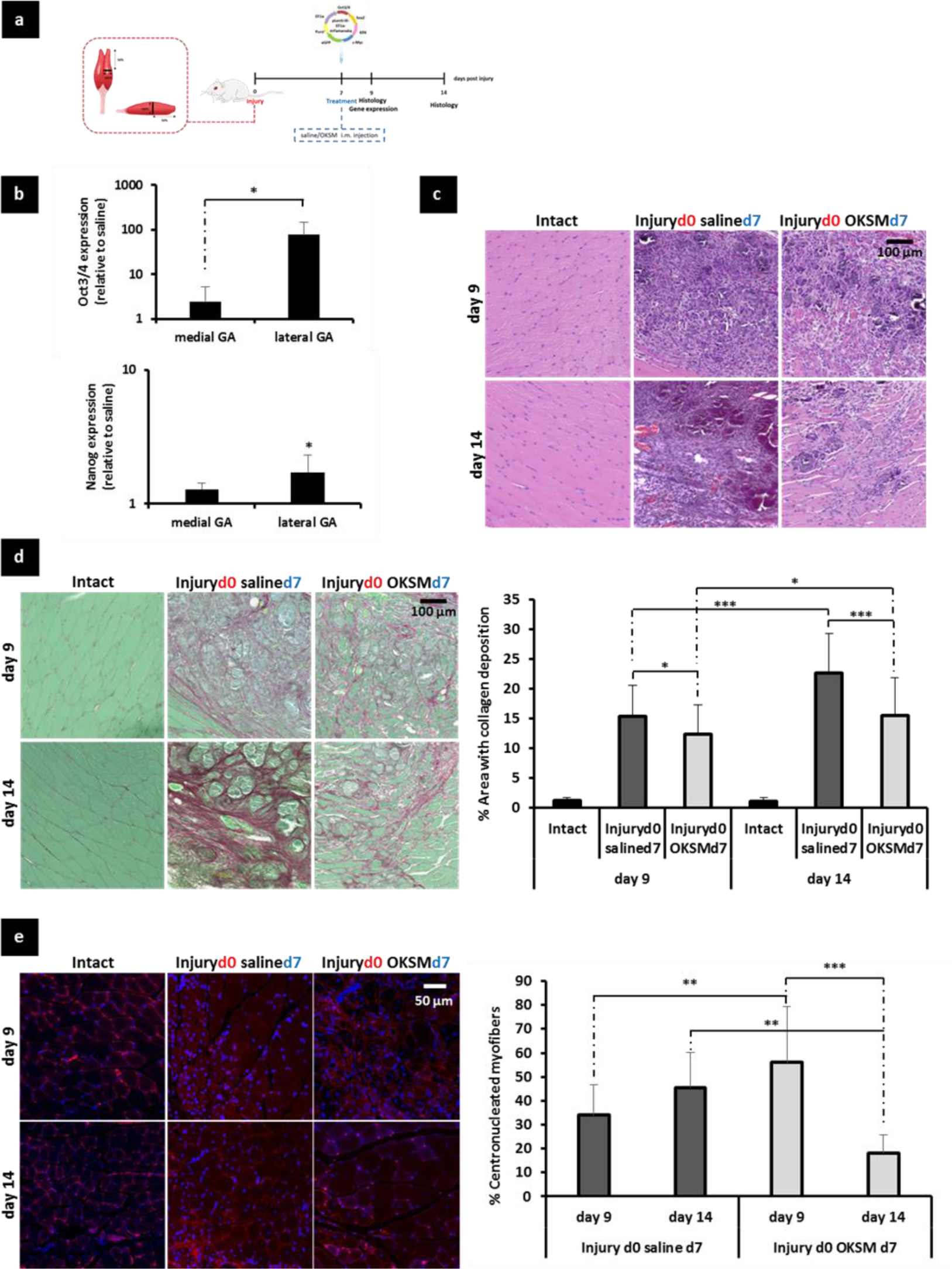
Effect of *in vivo* reprogramming towards pluripotency after laceration of the medial GA muscle. **(a)** The medial GA muscle of BALB/c mice was surgically lacerated and 100 μg pLenti-lll-EF1a-mYamanaka in 40 μl 0.9% saline or 40 μl saline alone were i.m. administered 7 days after the injury. GA muscles were dissected 9 and 14 days after injury. (**b**) *Oct3/4* and *Nanog* gene expression was normalised to the saline-injected group. *p<0.05 indicates statistically significant differences between *Oct3/4* expression in the medial and lateral GA muscle of pDNA injected mice and in the *Nanog* expression between the lateral GA muscle of pDNA and saline-injected animals, assessed by one-way ANOVA and Tukey’s test, n=4. (**c**) H&E staining (40X, scale bar represents 100μm). (**d**) Picrosirius red – fast green staining (40x, scale bar represents 100 μm) and measurement of areas with collagen deposition. *p<0.05 and ***p<0.001 indicate statistically significant differences between pDNA and saline-injected groups and different time points, assessed by one-way ANOVA and Tukey’s test (n=4 GA muscles per group, 2 sections per muscle, 5 random fields per section). (e) Laminin/DAPI staining (63X, scale bar represents 50 μm) and quantification of % centronucleated myofibers. **p<0.01 and ***p<0.001 indicate statistically significant differences between pDNA and saline-injected groups and different time points, assessed by Welch ANOVA and Games Howell’s test, (n=4 GA muscles per group, 2 sections per muscle, 3 random fields per section). All data are presented as mean ± SD.

H&E staining confirmed the presence of necrotic fibers and inflammatory infiltrates in the vicinity of the injury, which was evident in both groups on day 9, but diminished considerably by day 14 in the reprogrammed tissue only (Figure 5c). The formation of a collagen-based fibrotic scar is one of the hallmarks of muscle remodelling after injury that prevents the functional rehabilitation of the tissue [13, 14]. We investigated whether the administration of reprogramming factors would have any effect on the deposition of collagen after laceration and found that collagen-positive areas (detected by picrosirius red/fast green staining) on day 9 were more extensive in the tissues from saline-control animals, this difference being significantly more pronounced on day 14. Between these two time points, the fibrotic area (calculated as a percentage of the total area in the cross-section) in the vicinity of the injury increased only slightly (12.3 ± 5.1% to 15.5 ± 6.5%) in the OKSM-injected group. In contrast, the area of fibrotic tissue increased significantly, from 15.4 ± 5.2% on day 9 to 22.6 ± 6.7% on day 14 in the animals that received saline only (Figure 5d). Finally, laminin/DAPI staining allowed the quantification of centronucleated myofibers (Figure 5e). On day 9, a higher percentage of myofibers surrounding the injured site were regenerating in the reprogrammed group (56.0 ± 23.4%) – as evidenced by the centralised position of the nucleus – compared to the saline control (33.9 ± 12.9%). In the saline-injected group, the percentage of centronucleated myofibers continued to increase (45.6 ± 14.6% on day 14) and was considered an indication that the regeneration process was still ongoing, contrary to the *in vivo* reprogrammed tissue, in which the number of fibers regenerating decreased significantly (18.2 ± 7.7% on day 14). Collectively, these results suggested that *in vivo* reprogramming prevented the excessive deposition of collagen observed in injured muscles that did not receive reprogramming factors and accelerated the regeneration of the tissue at the histological level.

## Discussion

This study demonstrated that the induction of a transient pluripotent-like state within the skeletal muscle can accelerate tissue regeneration at the histological level in an injury model. Our laboratory was first to report *in vivo* cell reprogramming to pluripotency in adult mammalian tissue (liver) [3, 9]. Here, transient expression of the Yamanaka factors in healthy adult skeletal muscle also rapidly triggered the upregulation of pluripotency genes that are otherwise repressed in adult tissues. The expression of such markers persisted only briefly in the tissue and was accompanied by the upregulation of *Pax3*, a marker of myogenic precursors, and the downregulation of *MyoD1*, characteristically expressed at more mature stages in myogenesis (Figures 1 and 2). Altogether, a rapid and transient activation of an embryonic-like gene expression program was reproducibly observed in different WT and transgenic mouse strains and with the administration of different reprogramming pDNA.

The lack of contribution from satellite cell and pericyte proliferation to these observations was evidenced in a number of findings. First, we found that gene expression of several satellite- and pericyte-specific markers was downregulated in the pDNA-injected muscle tissues compared to saline controls (Figures 2e and 2f). In addition, *MyoD1* – which is rapidly upregulated upon satellite cell activation and highly expressed in their proliferating progeny [48] – was significantly downregulated in the reprogrammed specimens throughout our work (Figures 1d and 2d). The lack of satellite cell and pericyte involvement was reaffirmed in histological studies, in which we did not observe any differences in the abundance of such cells between the pDNA and saline-injected tissues (**Figure S1c**). Conversely, co-localisation studies in Nanog-GFP transgenics confirmed that pluripotent-like cells generated in the tissue upon *in vivo* reprogramming did not express satellite cell or pericyte markers. Finally, similar studies on a Pax3-GFP mouse strain confirmed that cells expressing the pluripotency marker NANOG, also expressed PAX3 (Figures 3 **and S1**), closer to the identity of early myogenic progenitors in developing skeletal muscle.

Intramuscular injection of pDNA has been previously reported to result in long-term transgene expression that can last for 2 months and reach peak levels 14 days after injection, based on the post-mitotic status of the myofibers[34]. This contrasted with the rapid decrease in the expression of the transfected reprogramming factors observed in our studies (Figures 1 and 2) and was explained by the proliferative status of the transfected cells evidenced by BrdU labelling (Figure 4d). No significant BrdU incorporation was observed in saline-injected controls and therefore proliferation was attributed to the reprogramming event. Indeed, active cell division is an early and mandatory step in the differentiated-to-pluripotent conversion[46]. This finding also suggested that *in vivo* reprogramming to a pluripotent-like state could enhance repopulation of the cell pool in the tissue following injury, precisely by increased cell proliferation. In addition, our histological observations of the evolution of the reprogrammed cell clusters over time suggested that, after the proliferative phase, reprogrammed cells re-differentiated and successfully integrated in the muscle tissue. This was supported by the gradual disappearance of mononucleated cell clusters, the rapid decrease in the expression of pluripotency markers, together with the emergence of desmin-positive centronucleated myofibers from days 4 to 8 after injection, absence of pronounced apoptosis and of dysplastic lesions or teratomas (Figures 4 **and S4**). Despite the consistency among these observations, specific lineage tracing studies that could allow permanent labelling of the *in vivo* reprogrammed cells should be pursued in the future to allow determination of their fate within the tissue and confirm the hypothesis that is postulated here. We also noted that centralised nuclei persisted only for up to 8 days after injection, which differs from classic regeneration where such feature is observed for longer periods of time [10].

Indeed, one of the caveats in our study remains the elucidation of the identity of those cells within the skeletal muscle tissue that were reprogrammed to a pluripotent-like state. Fully differentiated myofibers are known to uptake naked pDNA [29], making them interesting targets for gene delivery *in vivo.* However, their multinucleated and post-mitotic status would require cell fission and cell cycle re-entry to achieve successful reprogramming to pluripotency. While myofibers of urodele amphibians are able to de-differentiate and cleave into mononucleated progenitors during regeneration [50], there is currently no evidence that mammalian myofibers can undergo such processes. Recent studies have attempted to achieve mammalian myofiber de-differentiation and fission via ectopic expression of transcription factors that mediate such processes in the amphibian organism [51-53] or via muscle injury [54, 55]. However, these reports also lack robust lineage tracing tools that could confirm the origin of the resulting mononucleated cells. Myoblasts have been successfully de-differentiated into iPS cells *in vitro* [56] via the downregulation of *MyoD1*, orchestrated by the master regulator of pluripotency, *Oct4* [37]. Interestingly, we consistently observed a decrease in the mRNA levels of *MyoD1* in the reprogrammed groups throughout our study (Figures 1 and 2). We also confirmed downregulation of markers specific to satellite cells. This cell population has shown to be more efficiently reprogrammend than other more committed cell types [56]. Overall, and given the probably random incorporation of pDNA in a variety of cells, it is conceivable that different interstitial cells were reprogrammed rather than a specific cell type, all down-regulating their differentiated markers.

We have previously hypothesized that the generation of transiently proliferative, pluripotent-like cells in the context of an injured tissue might help its cellular repopulation and overall regeneration[33]. Indeed, there are notorious limitations in the management of major skeletal muscle injuries, for which the clinically-established treatment continues to be conservative (rest, ice, compression and elevation in the earliest stages, followed by physical therapy in the long term) and the often lack of complete recovery leads to fibrosis and loss of muscle contractility. In our study, the administration of pDNA encoding Yamanka factors 7 days after a clinically-relevant injury not only accelerated the clearance of necrosis and inflammatory infiltration (Figure 5c) and the appearance of centronucleated myofibers (Figure 5e), but also prevented the excessive collagen deposition observed in the control group (Figure 5d). While these results are encouraging, future studies are warranted to investigate whether such histological observations translate into significant differences in the recovery of muscle function and otherwise to identify the optimum gene transfer vectors and dosage regimes that may provide improved efficacy.

Other groups have explored the concept of *in vivo* reprogramming towards pluripotency in mammalian tissues following strategies by which the expression of Yamanaka factors was sustained over prolonged periods of time [4-7]. The fact that all such studies led to uncontrolled proliferation, random re-differentiation into multiple cellular lineages and remarkable tumorigenesis highlights the absolute necessity for transient reprogramming strategies, as the one described here and in our previous work in mouse liver [3, 8], to shape *in vivo* reprogramming towards pluripotency as a clinically-relevant alternative. Confirming this, Ohnishi et al. demonstrated that successful re-differentiation and re-integration of reprogrammed cells into their host tissues, which maintained their physiological functions intact, was feasible when the expression of reprogramming factors via a doxycycline-inducible promoter was limited to a maximum of 7 days. Conversely, the administration of doxycycline for longer periods of time resulted invariably in the generation of dysplastic lesions and tumorigenesis [5].

The teratoma-free approach that we have postulated in this work may also offer advantages to other reprogramming-based strategies. First, it bypasses the need for *ex vivo* manipulation and transplantation, and their associated risks, encountered by the *in vitro* generation of iPS cells. Such limitations have been exhaustively reviewed by others [59, 60]. The presence of a transient proliferative stage, evidenced in our studies by the occurrence of BrdU^+^ cell clusters (Figure 4d), offers a possibility for cellular expansion and repopulation that the *in vivo* transdifferentiation strategies described to date (i.e. direct reprogramming between two differentiated cell types, avoiding pluripotent intermediates and active cell division) lack[61-69]. In addition, transdifferentiation strategies require the identification of specific reprogramming factors for each particular cell type conversion, while reprogramming a variety of starting cell types towards the pluripotent state can be achieved with a defined cocktail of factors [1, 70, 71].

Likewise, this approach could also overcome some of the roadblocks faced by strategies that are currently being explored at the experimental level for the treatment of muscle laceration injuries. Among them, the administration of insulin-like growth factor-1 (IGF-1), basic fibroblast growth factor (bFGF) or nerve growth factor (NGF) has shown to improve muscle regeneration, but their clinical application suffers from their rapid clearance and, in some cases, controversial mitogen status[19]. Anti-fibrotic therapy has also been extensively researched through molecules that block the TGF1-β pathway, including suramin[16], gamma interferon[17], decorin[13], relaxin[18] and the combination of IGF-1 and decorin[14]. However, it has failed to achieve complete muscle rehabilitation due to lack of significant myogenesis. Cell therapy strategies have precisely aimed to address cellular repopulation of the injured site, but they are also not devoid of limitations. Initial efforts of myoblast transplantation, theoretically favoured due to their natural commitment for the myogenic lineage, were hampered by the extremely poor cell survival after transplantation[72]. Other cell types, including bone marrow mesenchymal stromal cells (BMMSCs)[20], CD133+ human peripheral blood cells[21], adipose tissue-derived regenerative cells (ADRCs)[22] and adipose tissue-derived stem cells (ADSCs)[24, 25] have reported very poor engraftment in the host tissue after laceration. Only Shi *et al.* were able to relate the improvement in muscle regeneration to successful integration, survival and differentiation of CD133^+^ peripheral blood cells to the myogenic and endothelial lineage[21], contrary to the rest of the studies all of which were mainly attributed to paracrine effects [22] [20]. Moreover, administration of the replacement cells at the time of injury significantly compromises clinical relevance in such studies, in combination with the extended culturing – sometimes close to 4 weeks – required to achieve sufficient cell numbers for implantation [20, 21, 24, 25]. In our approach, the Yamanaka factors were administered 7 days after injury prompted by the higher upregulation of *Nanog* achieved in comparison to their administration at the time of injury and 5 days later (**Figure S5b**). We reasoned that at earlier time points the presence of inflammatory infiltrates as well as myofibers undergoing injury-mediated necrosis would compromise the efficiency of transcription factor overexpression and reprogramming. We consider this finding particularly relevant to the clinical translation of *in vivo* reprogramming towards pluripotency, since it proves its efficacy even one week after injury, therefore circumventing the need for a rapid post-injury intervention.

Finally, another study has also explored a nucleic acid-based strategy for the regeneration of skeletal muscle after laceration. Nakasa *et al.* tested a cocktail of miRNAs involved in muscle development, which improved muscle regeneration at the histological and functional levels via the enhancement of myoblast proliferation, self-renewal and differentiation [26]. While promoting differentiation towards the myogenic lineage might be a more direct approach to repopulate injured muscle, avoiding pluripotent-like intermediates, it relies on the presence of sufficient numbers of muscle progenitors in the tissue, which may not be the case depending on the pathological condition to tackle. Muscle progenitors are known to be depleted as a consequence of ageing [73] and in some conditions such as muscular dystrophy, where muscle mass is progressively replaced by fibrotic and adipose tissue [74]. We advocate that *in vivo* reprogramming towards pluripotency can result in a more versatile tool to induce cellular repopulation in focal regions of injured tissue thanks to the reported universality of the Yamanaka factors to reprogram different cell types towards pluripotency [1, 70, 71].

We have proposed here a transient reprogramming strategy to generate pluripotent-like intermediates *in situ* that can assist cellular repopulation of an injured site, while circumventing the challenges faced by the transplantation of replacement cells - including cell isolation, extensive culturing and its inherent risk of genomic aberrations, delivery and engraftment. We hypothesize that this strategy could benefit from the presence of differentiation cues in the tissue microenvironment able to re-differentiate the *in vivo* reprogrammed cells to the appropriate phenotypes. The transient character of this approach has proved critical to avoid tumorigenesis, a burden that keeps other *in vivo* reprogramming to pluripotency strategies far from the road into the clinic.

## Methods

### DNA plasmids (pDNA)

Reprogramming plasmids pCX-OKS-2A encoding Oct3/4, Klf4, Sox2; pCX-cMyc encoding cMyc and pLenti-lll-EF1a-mYamanaka encoding Oct3/4, Klf4, Sox2, cMyc and eGFP were purchased as bacterial stabs from Addgene (USA) and Applied Biological Materials (USA), respectively. Research grade plasmid production was performed in Plasmid Factory, (Germany).

### Animals

All experiments were performed with prior approval from the UK Home Office under a project license (PPL 70/7763) and in strict compliance with the Guidance on the Operation of the Animals (Scientific Procedures) Act 1986. BALB/c mice were purchased from Harlan (UK). Sv129-Tg(Nanog-GFP) mice, which carry the eGFP reporter inserted into the *Nanog* locus [75], were a kind gift from the Wellcome Trust Centre for Stem Cell Research, University of Cambridge (UK) and bred and genotyped at the University of Manchester. CBA-Tg(Pax3-GFP), in which eGFP replaces the *Pax3* coding sequence of exon 1 [76], were bred and genotyped at the University of Manchester. All the mice used in this work were female used at 7 weeks of age. Mice were allowed one week to acclimatize to the animal facilities prior to any procedure.

### Intramuscular (i.m.) administration of pDNA

Mice were anesthetised with isofluorane and the left gastrocnemius (GA) muscle was injected with either 50 μg pLenti-lll-EF1a-mYamanaka or 50 μg pCX-OKS-2A and 50 μg pCX-cMyc in 50 μl 0.9% saline solution. The contralateral (right) leg was injected with 50 μl 0.9% saline solution alone as internal control. For the dose-escalation study mice were injected with 100 μg pLenti-lll-EF1a-mYamanaka. Mice were culled at different time points, including 2, 4, 8, 12, 24, 50 and 120 days after i.m. injection, as specified in each particular study.

### RNA isolation and real-time qRT-PCR analysis

Aurum Fatty and Fibrous Kit (Bio-rad, UK) was used to isolate total RNA from muscle tissue. cDNA synthesis from 1 μg of RNA sample was performed with iScript cDNA synthesis kit (BioRad, UK) according to manufacturer’s instructions. 2 μl of each cDNA sample were used to perform real-time RT-qPCR reactions with iQ SYBR Green Supermix (Bio-Rad, UK). Primer sequences are shown in **Supplementary Table 1**. Experimental duplicates were run on CFX-96 Real Time System (Bio-Rad, UK) with the following protocol: 95°C for 3 min, 1 cycle; 95°C for 10 sec, 60°C for 30 sec, – repeated for 40 cycles. *β-actin* was used as a housekeeping gene and gene expression levels were normalised to saline-injected controls.

### Immunohistochemistry (IHC) of BALB/c muscle tissue sections

GA muscles were dissected 2 and 4 days after i.m. injection with pLenti-lll-EF1a-mYamanaka or saline control and immediately frozen in isopentane, pre-cooled in liquid nitrogen. 10 μm-thick sections (in the transverse plane) were prepared on a cryostat (Leica, CM3050S) and air-dried for 1 h at room temperature (RT) prior to storage at −80°C. Before staining, muscle sections were post-fixed with methanol, pre-cooled at −20°C, for 10 min, air-dried for 15 min and finally washed twice with PBS for 5 min. After washing, the sections were incubated for 1 h in blocking buffer (5% goat serum-0.1% Triton in PBS pH 7.3) at RT, followed by two washing steps with PBS (1 %BSA- 0.1% Triton, pH 7.3) before overnight incubation at +4°C with the different primary antibodies: rabbit pAb anti-OCT4 (ab19857,3 μg/ml, Abcam, UK), rabbit pAb anti-SOX2 (ab97959, 1 μg/ml, Abcam, Uk) and rabbit pAb anti-NANOG (ab80892, 1 μg/ml, Abcam, UK), Next day sections were washed (2 min each) with PBS and incubated (1.5 h at RT) with the secondary antibody (goat polyclonal anti-rabbit IgG labeled with Cy3, 1/250, Jackson ImmunoResearch Laboratories Inc.). After two washes in PbS and mounting in ProLong^®^ Gold anti-fade DAPI containing mountant (Life Technologies, UK). For GFP signal (from the reporter in pLenti-lll-EF1a-mYamanaka) tissue sections were directly mounted in ProLong^®^ Gold medium containing DAPI after fixation. 40X images were collected on a Zeiss Axio Observer epi-Fluorescence microscope.

### Alkaline phosphatase (AP) staining of muscle sections

AP activity staining was performed using the BCIP/NBT liquid substrate system (Sigma, UK). Frozen muscle sections were prepared as described above and incubated with BCIP/NBT liquid substrate system for 30 min. Sections were washed with water and mounted with glycerol gelatin aqueous mounting media (Sigma, UK). Random images (10X) were captured by light microscopy.

### Characterisation of GFP^+^ cell clusters in Nanog-GFP and Pax3-GFP transgenic mice

To characterize the clusters of reprogrammed cells in Nanog-GFP and Pax3-GFP mice, GA muscles were administered with pCX-OKS-2A and p-CX-cMyc pDNA vectors to avoid overlapping of the eGFP reporter in pLenti-lll-EF1a-mYamanaka and sacrificed 2 and 4 days after injection. To observe the GFP signal frozen tissue sections obtained as described before were simply mounted with ProLong^®^ Gold anti-fade DAPI containing mountant (Life Technologies, UK) after fixation and imaged with 3D Histech Pannoramic 250 Flash slide scanner and Panoramic Viewer Software (100X). To investigate co-localisation of the GFP signal with the expression of different markers, tissue sections were processed for IHC as described before. Rabbit pAb anti-OCT4 (ab19857,3 μg/ml, Abcam, UK), rabbit pAb anti-NANOG (ab80892, 1 μg/ml, Abcam, UK), rabbit pAb anti-AP (ab95462, 1:200, Abcam, UK), mouse mAb anti-SSEA1 (ab16285, 20 μg/ml, Abcam, UK), rabbit pAb anti-Pax7 (pab0435, 1:200, Covalab, France), and rabbit mAb anti-PDGFrβ (#3169, 1:100, Cell Signalling) were used as primary antibodies. Anti-PDGFrβ was a kind gift from Prof. Cossu’s lab (University of Manchester). Goat pAb anti-rabbit IgG labeled with Cy3 and goat pAb anti-mouse IgG labeled with Cy5 (1/250, Jackson ImmunoResearch Laboratories Inc) were used as secondary antibodies. Images were taken at 100X magnification with a Leica TCS SP5 AOBS inverted confocal microscope.

### Histological evaluation of muscle tissue

The GA muscles of BALB/c mice were dissected on days 2, 4, 8, 15, 50 and 120 after i.m. injection with pLenti-lll-EF1a-mYamanaka or saline control, frozen and sectioned as previously described and H&E following a standard protocol. Tissue sections were imaged with a 3D Histech Pannoramic 250 Flash slide scanner and representative images at 40X and 100X magnification were taken with Pannoramic Viewer software.

### Desmin/laminin/DAPI staining

10 μm-thick cryosections were stained following the IHC protocol previously described using rabbit pAb anti-laminin (ab11575, 1:200, Abcam, UK) and mouse mAb anti-desmin (ab6322, 1:200, Abcam, UK) primary antibodies and goat pAb anti-rabbit IgG labeled with Cy3 (1/250, Jackson ImmunoResearch Laboratories Inc) and goat pAb anti-mouse IgG labeled with Cy5 (1/250, Jackson ImmunoResearch Laboratories Inc) as secondary antibodies. 40X images were obtained with a Zeiss Axio Observer epi-Fluorescence microscope.

### BrdU labelling and detection of proliferating cells

BrdU assay was used to label proliferating cells *in vivo*, as previously described[77]. BALB/c mice were intraperitoneally (i.p.) with 500mg/kg of 5-Bromo-2’-deoxyuridine (BrdU, B5002, Sigma, UK) in 0.9% saline 18 h after i.m. injection with pLenti-lll-EF1a-mYamanaka or saline control. 6 h later, GA muscles were dissected and processed for IHC as described above. Treatment with 2 N HCl (10 min at 37°C) after fixation of the sections was performed to denature the DNA. Mouse mAb anti-BrdU (B8434, 1:100, Sigma, UK) and goat pAb anti-mouse IgG labeled with Cy5 (1/250, Jackson ImmunoResearch Laboratories Inc) were used.100X images were captured with a Leica TCS SP5 AOBS inverted confocal microscope.

### Surgically-induced skeletal muscle injury model and *in vivo* reprogramming to pluripotency

BALB/c mice were anesthetised with isoflurane and the left hind limb was shaved and prepared for surgery. 0.05 mg/kg buprenorphine was subcutaneously (s.c.) administered at the start of the intervention. A vertical skin incision (6 mm long) was made overlying the posterior compartment of the calf with a scalpel number 11 and the fascia was exposed and incised with fine scissors at the level of mid-gastrocnemius to release the muscle belly. The medial gastrocnemius muscle was blunt dissected and transected at its mid-point with a single incision in the transverse plane, as described in **Figure S6**, preserving the sural nerve and the lateral gastrocnemius. The skin was sutured with 5 interrupted stitches (Vicryl 6-0 absorbable suture, Ethicon, UK) and the mice were allowed to recover in a warm chamber. The contralateral (right) leg was left intact for internal control. 7 days after the surgery 100 μg pLenti-lll-EF1a-mYamanaka in 40 μl 0.9% saline or 40 μl 0.9% saline alone were i.m. administered in the injured (left) leg. The contralateral (right) leg was left uninjected. Animals were sacrificed at days 9 and 14 after the injury (2 and 7 days after the administration of pDNA, respectively) for histological and electromechanical investigations (n=4). A group of animals was also culled at day 9 (day 2 after i.m. injection) for gene expression analysis.

### Laminin/DAPI staining and analysis of % of centronucleated myofibers

10 μm-thick cryosections were obtained 7 days after CTX administration and 9 and 14 days after surgical laceration and IHC stained following the protocol previously described using rabbit pAb anti-laminin (ab11575, 1:200, Abcam, UK) and goat pAb anti-rabbit IgG labeled with Cy3 (1/250, Jackson ImmunoResearch Laboratories Inc) antibodies. 10X images were collected using a Zeiss Axio Observer epi-Fluorescence microscope (CTX study) or a 3D Histech Pannoramic 250 Flash slide scanner and Pannoramic Viewer Software (laceration study). The number of centronucleated myofibers and total number of fibers per cross-sectional area were quantified using ImageJ 1.48 software from 2 to 3 randomly selected fields per section. The data for each mouse were calculated from 2 sections and we observed 3 to 4 mice in each group, as specified in each study. The measurements and calculations were conducted in a blinded manner.

### Picrosirius red-fast green staining and analysis of fibrotic area

GA muscles were dissected 9 and 14 days after laceration injury and fixed in 10% buffered formalin solution (Sigma, UK). Tissues were embedded in paraffin wax and 5 μm thick sagittal and transverse sections were obtained with a microtome (Leica RM2255) and left to dry overnight at 37°C prior to the staining procedure. Tissue sections were de-paraffinized following a standard protocol and stained for 1 h in picrosirius red/fast green staining solution (0.1% Sirius red, 0.1% fast green in a saturated aqueous solution of picric acid). The tissue sections were then quickly immersed in 0.5% acetic acid for 6 s, dehydrated in 100% ethanol and xylene to be finally mounted with DPX mountant (Sigma, UK). Sections were imaged on a 3D Histech Pannoramic 250 Flash slide scanner and 40X images were obtained with Pannoramic Viewer Software. The percentage of collagen-positive areas was quantified with ImageJ 1.48 software in 5 randomly selected fields per section. The data for each mouse were calculated from 2 sections and 4 mice were included per group. The measurements and calculations were conducted in a blinded manner.

### Statistical analysis

N numbers were specified for each particular study. Statistical analysis was performed first by Levene’s test to assess homogeneity of variance. When no significant differences were found in the variances of the different groups, statistical analysis was continued by one-way ANOVA and Tukey's post-hoc test. When variances were unequal, the analysis was followed with Welch ANOVA and Games-Howell's post-hoc test. Probability values <0.05 were regarded as significant. SPSS software version 20.0 was used to perform this analysis.

## Acknowledgements

Irene de Lázaro would like to thank Obra Social LaCaixa and UCL School of Pharmacy for a jointly funded PhD Studentship. The authors would also like to acknowledge the Histology and Bioimaging Facilities at the University of Manchester for assistance, as well as Dr Adam Reid (University of Manchester) for his help in the establishment of the injury model utilised in this study.

## Author contributions

IdL, AY and KK conceived the project and designed experiments. IdL and AY performed the majority of experiments. SQ contributed to some qPCR data. YN and FR performed some IHC experiments. GC advised in experimental design and interpretation of data. IdL, AY, GC and KK wrote the manuscript.

